# Histamine H_1_-Receptor-Mediated Modulation of Nmda Receptors

**DOI:** 10.1101/2021.11.02.466417

**Authors:** J-M. Arrang, V. Armand

**Author notes:** Corresponding author: Vincent Armand, Université de Paris, SPPIN – Saints-Pères Paris Institute for the Neurosciences, CNRS, Paris F-75006, France Descartes 45, rue des Saints Pères 75006 Paris, France. The authors contibuted equally to the study.

## Abstract

This study attempted to clarify the role of histamine H_1_ receptors in epilepsy by exploring the effects of agonists and inverse agonists on the rundown of the current induced by iterative applications of NMDA or GABA. Mepyramine, a classical H_1_-receptor antagonist / inverse agonist, increased the NMDA current by about 40 % during the first minutes of recording. This effect was concentration-dependent, maximal at 10 nM, and mimicked by triprolidine, another antagonist / inverse agonist. No endogenous histamine being detected in the cultures by a selective immunoassay, both compounds were acting as inverse agonists. Indicating a high constitutive activity of the H_1_ receptor in this system, histamine did not affect on the NMDA rundown, including its settlement, but reversed significantly the effect of mepyramine. A similar pattern was obtained with 2,3 bromophenyl histamine, a selective H_1_-receptor agonist. The initial increase induced by the two inverse agonists was followed by the same rundown as in controls. H_1_– and NMDA receptors colocalised in most cultured neuronal cells. Mepyramine and histamine did not affect on the GABA rundown. Our findings suggest an interaction between H_1_– and NMDA receptors. Inactivation of the H_1_-receptor by its inverse agonists delays the settlement of the NMDA rundown, which may underlie their proconvulsant effect reported in clinics. Therefore, H_1_-receptor constitutive activity, and the effect of histamine revealed in its absence, tend to facilitate the initiation of the rundown, which is consistent with the anticonvulsant properties of histamine via activation of H_1_-receptors reported in many studies.

## 1 INTRODUCTION

The effects of histamine in brain neurotransmission are mediated by three histamine receptor subtypes (H_1_, H_2,_ and H_3_), which are all G protein-coupled receptors. Histamine has long been known to decrease epileptogenic activity. This modulatory effect has been pharmacologically characterized in various animal models of epilepsy and is mediated by activation of H_1_ receptors. Activation of H_1_ receptors by endogenous histamine inhibits amygdaloid-kindled seizures in rats (Kamei 2001; Hirai *et al*, 2004), protects the developing rat hippocampus from kainic acid-induced neuronal damage (Kukko-Lukjanov *et al* 2006), protects rats from pentylenetetrazol-induced seizures (Zhang *et al* 2003; Nishida *et al* 2007), inhibits the development of seizures in the EL mouse, an animal model for hereditary temporal lobe epilepsy (Yawata *et al* 2004). Consistent with these findings, the proconvulsant effect observed after blockade of H_1_ receptors by brain penetrating antagonists, or after deletion of its gene in H_1_-receptor knock-out mice, is also well documented. First-generation H_1_ antagonists elicit epileptogenic activity in amygdala-kindled rats (Fujii *et al* 2003), cause behavioral and EEG seizures in rats (Kamei *et al* 2000), increase the duration of electrically induced seizures in developing mice (Yokoyama *et* la 1993). In H_1_-receptor knock out mice, the development of pentylenetetrazol-induced kindling (Chen *et al* 2003), or amygdaloid kindling (Hirai *et al*, 2004), is accelerated, and both intensity and duration of kainic acid-induced seizures are enhanced (Kukko-Lukjanov *et al* 2010). Last, the first generation of H_1_ antagonists, largely used as classical anti-allergy drugs, occasionally induced convulsions in healthy children and patients with epilepsy (Churchill 1949; Yokoyama & Iinuma 1996; Yasuhara *et al* 1998). If the involvement of brain H_1_ receptors in the anticonvulsant effect of histamine is well established, the molecular mechanisms leading to this anticonvulsant effect remain largely unknown. Epilepsy is traditionally thought to result from an imbalance of excitatory glutamate neurons and inhibitory GABAergic neurons, with an over-activation of the NMDA receptor and a hypoactivity of GABA receptors, both leading to the generation and spread of paroxysmal activities. In this study, we used the model of the rundown of the currents, observed both with NMDA and GABA receptors and the rundown results from different mechanisms, i.e., a dephosphorylation of the GABA receptor (Laschet *et al* 2007) and a calcium and ATP dependent modulation of the NMDA receptor (Rosenmund & Westbrook 1993). Previous studies have shown that the rundown of GABA currents is modified in temporal lobe epilepsy (Palma *et al* 2007) and in human partial epilepsy (Laschet *et al* 2007). The effect of the drugs was explored in rundown conditions, by applications separated by short-lasting intervals of the agonists NMDA or GABA, the iterative activation of receptors produces a time-dependent decrease in the currents we have explored the putative modulation of NMDA and GABA currents by histamine as well as high selective ligands for H_1_ receptors.

## 2 MATERIALS AND METHODS

### 2.1 Ethical approval

Animal euthanasia procedures were carried out in accordance with the Animal Protection Association of ethical standards and the French legislation concerning animal experimentation (authorization no. **75-1270**)

### 2.2 Primary neuronal cultures

Primary neuronal cultures from rat hippocampus were performed as previously described (Burban *et al* 2010). The hippocampus was removed from 18-day-old embryos of male Wistar rats (Janvier, Le Genest-St-Isle, France). Cells were dissociated and seeded on 12-well culture dishes (100, 000 cells/well containing 1 ml of medium), previously coated with poly D-lysine. After removal of the coating solution, cells were seeded in neurobasal medium supplemented with B27 (1:50), GlutaMAX-I– (2 mM) and penicillin–streptomycin (5 IU/ml and 5 µg/ml, respectively). In these conditions, cultures of neurons were favored neuron cultures to the detriment of glial cells. Neurons were maintained for 2 to 4 weeks at 37°C in a humidified atmosphere containing 5% CO2.

### 2.3 Whole-cell patch clamp

Ionic currents were recorded within large pyramidal neurons. For NMDA experiments, the external solution for recording whole-cell currents contained 140 mM NaCl, 5 mM KCl, 1.2 mM CaCl2, 12 mM HEPES acid, 12 mM HEPES sodium, and 33 mM D-glucose. Tetrodotoxin (1 µM) was added to eliminate the voltage-gated sodium channel currents. Glycine 5 µM was added to the external solution. The pH was 7.3 and the osmolarity was adjusted to 300 mOsm with D-glucose. The internal pipette solution contained 100 mM CsF, 40 mM CsCl, 1 mM CaCl2, 20 mM HEPES acid and 10 mM EGTA. The pH was adjusted to 7.3 with CsOH. A whole-cell amplifier (AXOPATCH 1D) was used to measure the current responses, with the membrane potential clamped at –60 mV. During the recording, NMDA (50 µM) was applied for 2 seconds every 30 seconds for 30 minutes.

For GABA experiments, the external solution for recording whole-cell currents contained 135 mM NaCl, 3 mM KCl, 2 mM CaCl2, 10 mM HEPES acid, 1 mM MgCl2, 7 mM triethylamine chloride and 10 mM D-glucose. Tetrodotoxin (1 µM) was added to eliminate the voltage-gated sodium channel currents. The pH was 7.3 and the osmolarity was adjusted to 300 mOsm with D-glucose. The internal pipette solution contained 130 mM CsF, 10 mM CsCl, 4 mM NaCl, 0.5 mM CaCl2, 10 mM HEPES acid, 5 mM EGTA and 7 mM Mg-ATP. The pH was adjusted to 7.3 with CsOH. A whole-cell amplifier (AXOPATCH 1D) was used to measure the current responses.

The membrane potential was clamped at –80 mV so that GABA application evoked inward current. During the recording, GABA (100 µM) was applied for 1 second every 3 minutes for 30 minutes (Laschet *et al* 2007). A rapid perfusion system was used for drug applications. Drugs were added both in bath and perfusion solutions.

During the patch clamp experiments the series resistance was monitored, cell with a > 30 % change during the 30 minutes of experiments were discarded from the analysis.

### 2.4 Histamine levels in cultures

Histamine levels in the culture medium were evaluated with an enzyme immunoassay (Immunotech) with a detection limit of 0,1 nM.

### 2.5 Immunohistochemistry of primary cultures

Primary cultures of hippocampal neurons were fixed with 2 % paraformaldehyde in phosphate buffer (0,1M pH 7,4) at room temperature for 10 min, rinsed for 48 h in 0.1M PBS phosphate buffer, primary cultures were double-immunolabeled for GABAAR/H1R; NMDAR/H1R; HUC/D/H1R.

Antibodies (dilution) used for immunofluorescent labelling were as follows: Anti-alpha 1 subunit of GABA_A_ receptor (1:1000) (NeuroMab); anti-NR1 subunit of NMDA receptor (1:1000) (Synaptic Systems – SYSY); anti-H_1_ receptor (1:100) (Alomone Labs – AL); and anti-HUC/D (pan-neuronal marker) (1:30) ( Invitrogen).

### 2.6 Drugs

NMDA, glycine, GABA, mepyramine, triprolidine and histamine were obtained from Sigma Aldrich (St Louis, MO USA). 2,3 Bromophenylhistamine dimaleate was provided by W. Schunack (Freie Universität Berlin, Germany).

### 2.7 Analysis of Data

Currents were normalized to the amplitude of the current recorded at t0 (first NMDA application, defined as 100%). The time-dependent variation of the current obtained in each condition (NMDA + drug) was compared to the control (NMDA alone) by a two-way ANOVA with Bonferroni adjustment. Statistical analysis was performed using a two-way ANOVA analysis for repeated measures followed by Dunnett’s post hoc analysis (one-tailed distribution). The data are expressed as percentages of control values +/− SD.

The area under the curve for each condition was calculated using cumulative values and expressed as a percent of control conditions. In this case, statistical analysis was performed using a Dunnett’s test since the samples were of a normal distribution.

## 3 RESULTS

### 3.1 Effect of mepyramine on the rundown of NMDA currents

In whole cells recordings from cultured hippocampal neurons, the successive application of the same concentration (50 µM) of NMDA at short intervals (every 30 s) was accompanied by a progressive decrease of the response, i.e., by a rundown of NMDA currents (Fig 1A). The amplitude of this rundown was similar to that previously reported (Rosenmund & Westbrook 1993). The H_1_– receptor antagonist / inverse agonist mepyramine used at 10 nM induced a progressive increase of the current during the first 10 min. Then, in a second phase, the rundown of currents was observed with a slope similar to that observed during the rundown in control conditions (Fig 1A, B). The area under the curve (AUC) generated during the 30 min of recordings showed that the increase induced by mepyramine during the first phase was concentration-dependent and saturable. The increase induced by mepyramine was already significant at 2 nM (14 ±4,1 %), with the maximal amplitude reached at 10 nM (39 ±4,1 % compared to 37 ± 2,8 % at 50 nM)(Fig4A). From these data, a half-maximal concentration of ∼3 nM could be estimated with a maximal effect of ∼40 % (Fig1C, Fig 4A).

**Figure 1.**
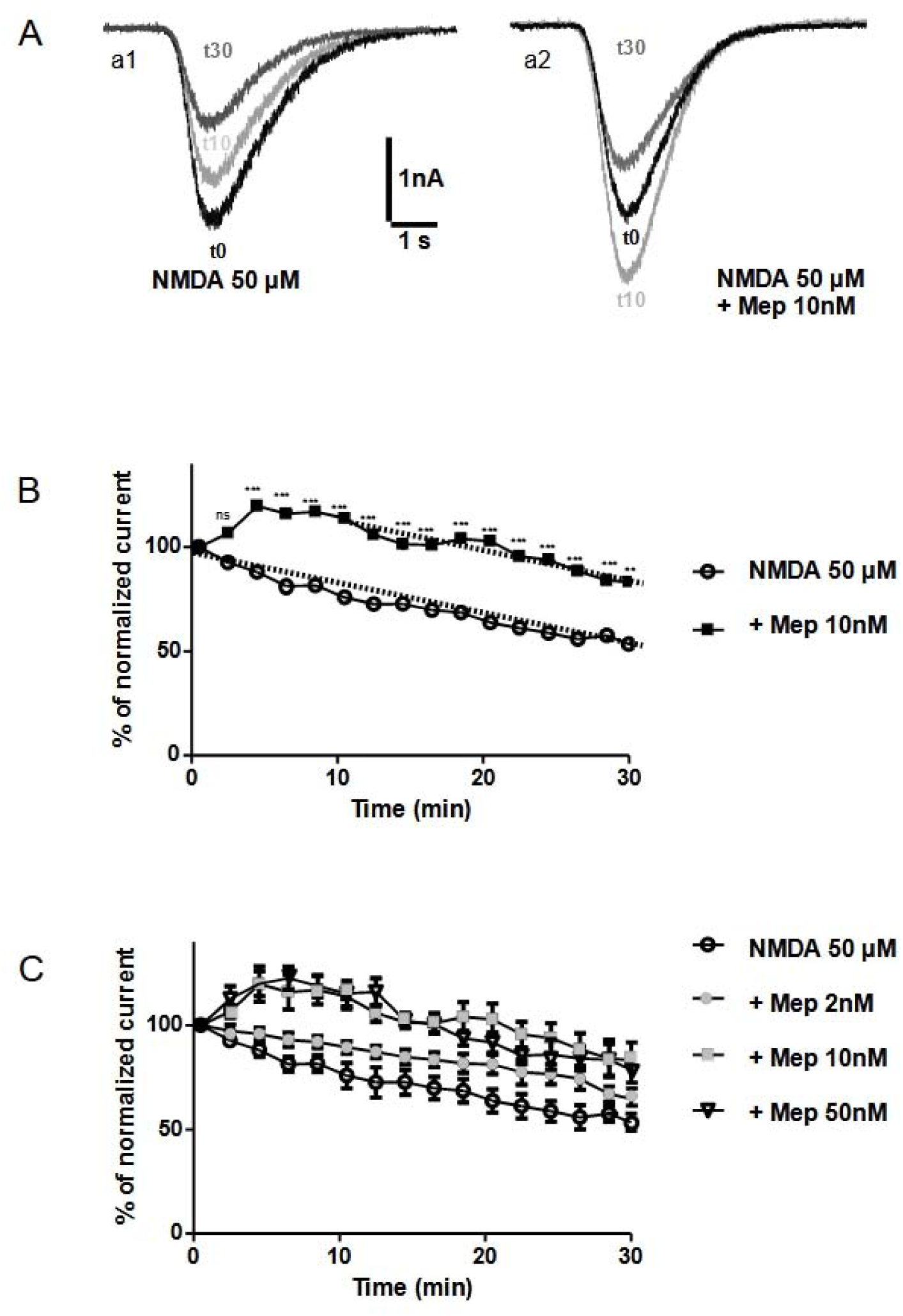
Effect of mepyramine (Mep) on the rundown of NMDA currents. Rundown of NMDA currents was observed by whole-cell recording. Inward currents were evoked by 2 s application of 50 µM NMDA at 30 s intervals. **A** Currents evoked after the first application (t0), after 10 min (t10) and 30 min (t30) of recording, are superposed. NMDA was applied alone (a1), or in the presence of mepyramine (a2). **B** Currents, normalized to the first application (t0=100%), were evoked by application of NMDA alone (n=10) or with mepyramine (n=10). The slopes of the rundown obtained in each condition remained unchanged, as shown by the corresponding dotted lines. ns: nonsignificant; **p<0.01; ***p< 0.001, vs NMDA alone. Note the error bars are in fig 1C. **C** The normalized currents evoked by NMDA alone (n= 10), and in the presence of 2, 10 and 50 nM mepyramine (n=10 for each condition), indicate that the effect of mepyramine is concentration dependent.

To go further on the increase induced by mepyramine, we explored a putative direct effect of the drug on the NMDA receptors. NMDA (50 µM) was applied three times at longer intervals (3 min) to avoid the settlement of any rundown. No difference could be found between the currents observed with and without mepyramine (n=10, data not shown), thereby excluding a direct effect of the drug with NMDA receptors.

### 3.2 Effect of triprolidine on the rundown of NMDA currents

Triprolidine, another classical H_1_ antagonist/inverse agonist, mimicked the effect of mepyramine and also delayed the rundown of NMDA currents during the first minutes of recording (Fig 2). Analysis of the AUC led to a significant increase compared to NMDA alone. However, the effect of triprolidine used at the same concentration (10 nM) was lower than that of mepyramine (25 ± 4,5 %) (Fig 4A).

**Figure 2.**
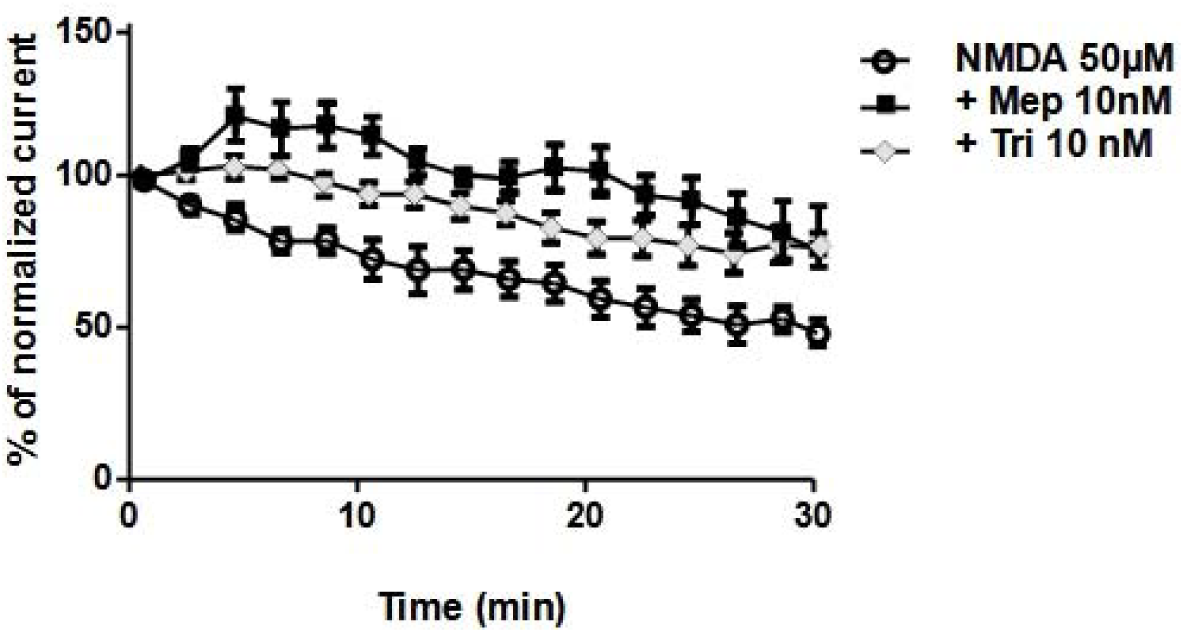
Effect of triprolidine (Tri) on the rundown of NMDA currents. The normalized currents were analysed after application of triprolidine (n=10), and compared to those obtained with NMDA alone or with mepyramine 10nM.

### 3.3 Effect of H_1_ agonists used alone on the NMDA rundown

The presence in the culture medium of endogenous histamine could not be detected by a selective immunoassay, indicating that mepyramine and triprolidine were acting as inverse agonists and not as antagonists against endogenous histamine. Exogenous histamine used at a high concentration (100 µM) had no effect on the NMDA rundown (Fig 3A). 2,3 Bromophenylhistamine, a more selective and potent H_1_ agonist, had also no effect on the NMDA rundown (Fig 3B).

**Figure 3.**
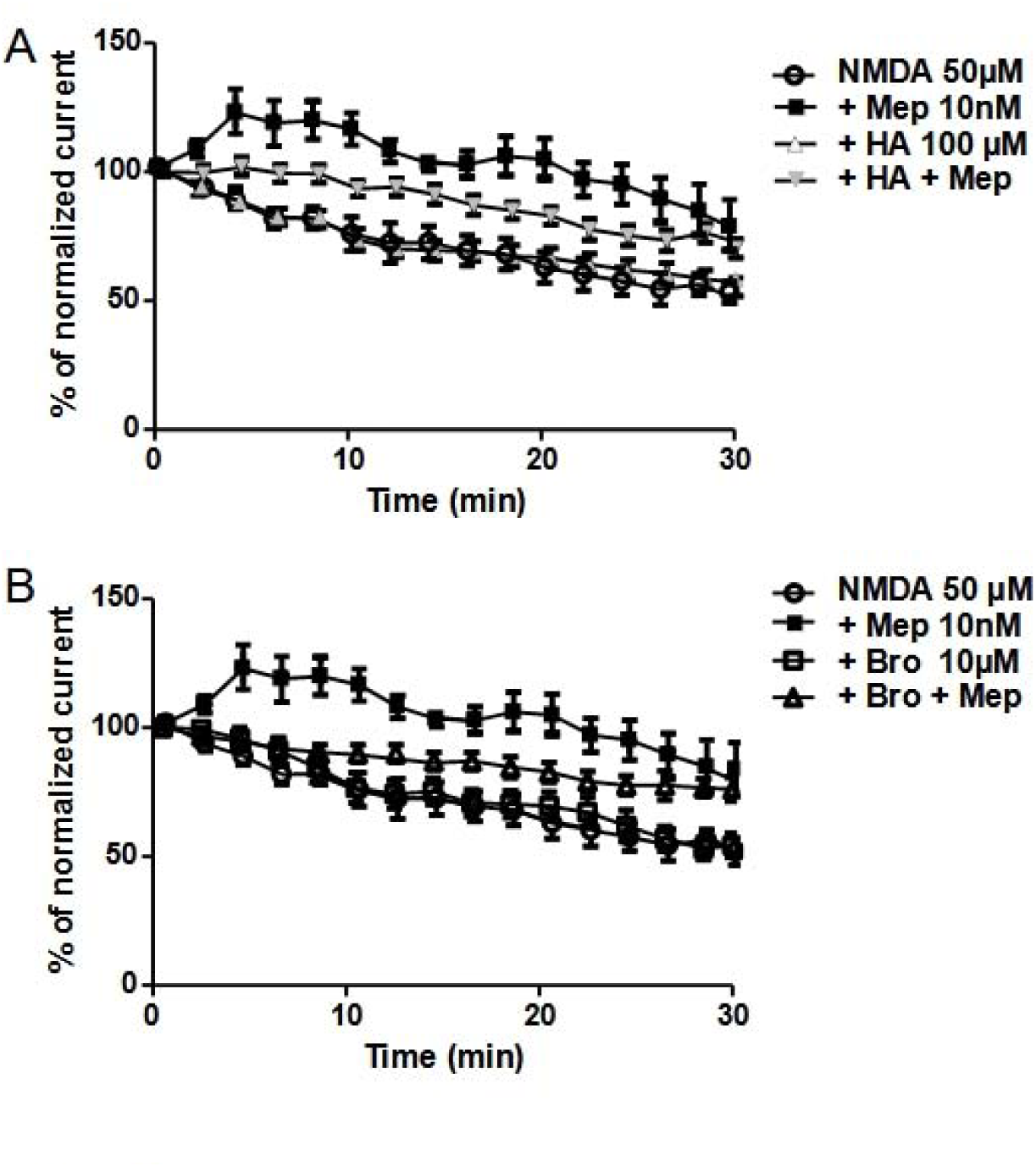
Effect of histamine (HA) and 2,3 bromophenylhistamine (Bro) on the rundown of NMDA currents compared to those obtained with NMDA alone or with mepyramine 10nM. **A** Effect of histamine alone (n=10) or with mepyramine (n=10) on the normalized currents evoked by NMDA. **B** Effect of 2,3 bromophenylhistamine (Bro) alone (n=10) or with mepyramine (n=10) on the normalized currents evoked by NMDA.

### 3.4 Effect of H1 agonists on the increase of currents induced by mepyramine

Agonists displaying no effect, and antagonists behaving in fact as inverse agonists on the response, we further explored the effects of agonists against mepyramine. Histamine (100 µM) significantly reversed by about 50 % the effect of the mepyramine (Fig 3A, Fig 4B). A similar profile was observed with 2,3 bromophenylhistamine used at a ten-fold lower concentration (10 µM). It also reversed, in a significant manner and with an even higher amplitude (by about 60 %), the effect of mepyramine (Fig 3B, Fig 4B).

**Figure 4.**
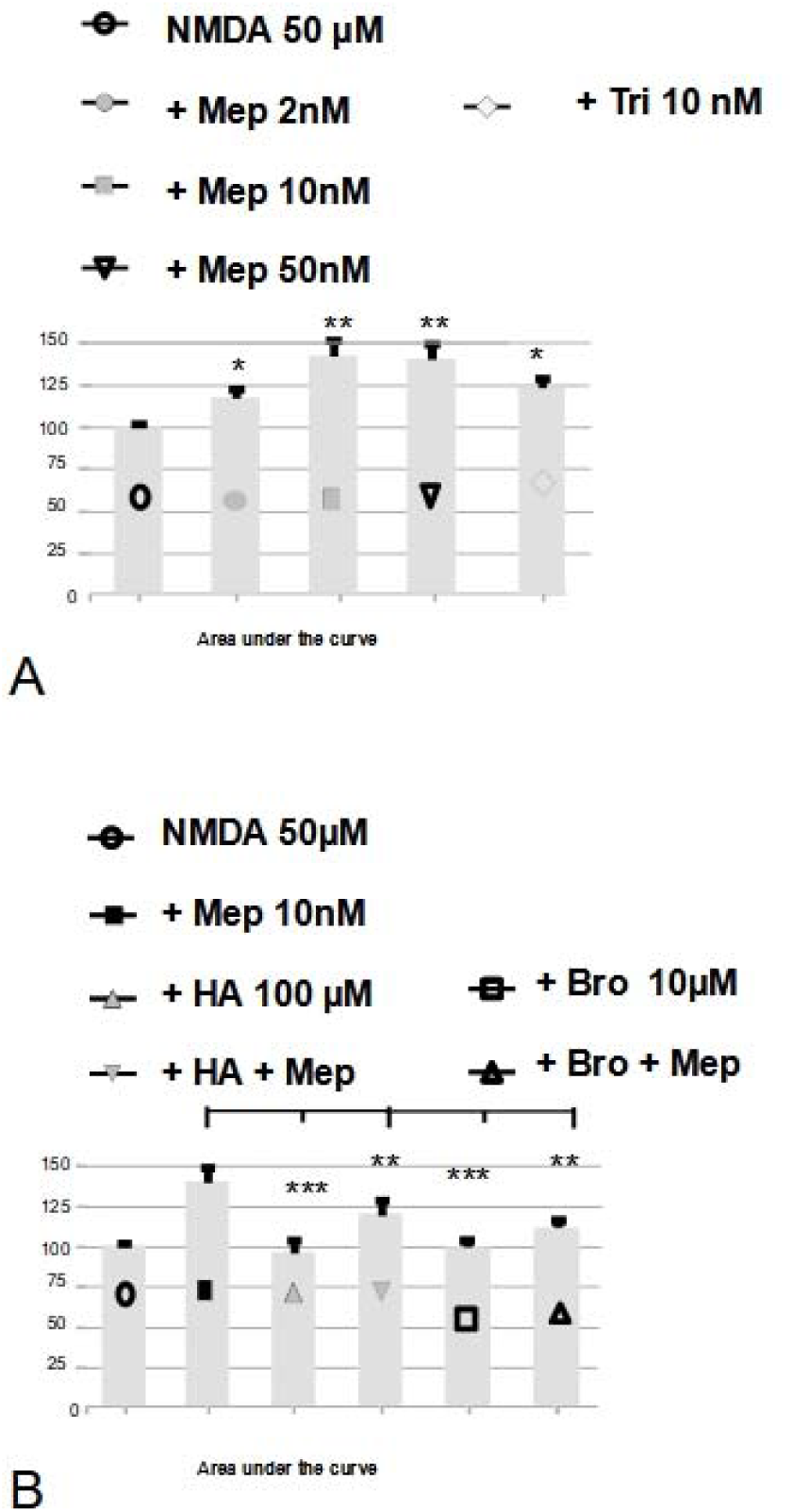
The area under the curve (AUC) for each condition was expressed as percent of the control (NMDA alone=100%). **A** The AUC were analysed after application of mepyramine (Mep) (2,10,50 nM) and triprolidine (Tri) (10nM), and compared to those obtained with NMDA. *p<0.05; **p<0.01 vs NMDA alone. **B** The AUC were analysed after application of histamine (HA) (100µM) and 2,3 bromophenylhistamine (Bro) alone (10µM) or added with mepyramine (10nM), and compared to those obtained with mepyramine alone (10 nM). **p<0.01, ***p<0.001 vs mepyramine (10nM).

### 3.5 Effect of histamine and mepyramine on the GABA rundown

In whole cells recordings from cultured hippocampal neurons, the expected rundown of the GABA_A_receptor was observed after iterative applications of 50 µM GABA (Fig 5A). The amplitude of the rundown was similar to that previously reported (Laschet *et al* 2007). Neither mepyramine used at 10 nM (Fig 5B) or 50 nM (not shown), nor histamine 100 µM (Fig 5B), had any effect on the GABA rundown.

**Figure 5.**
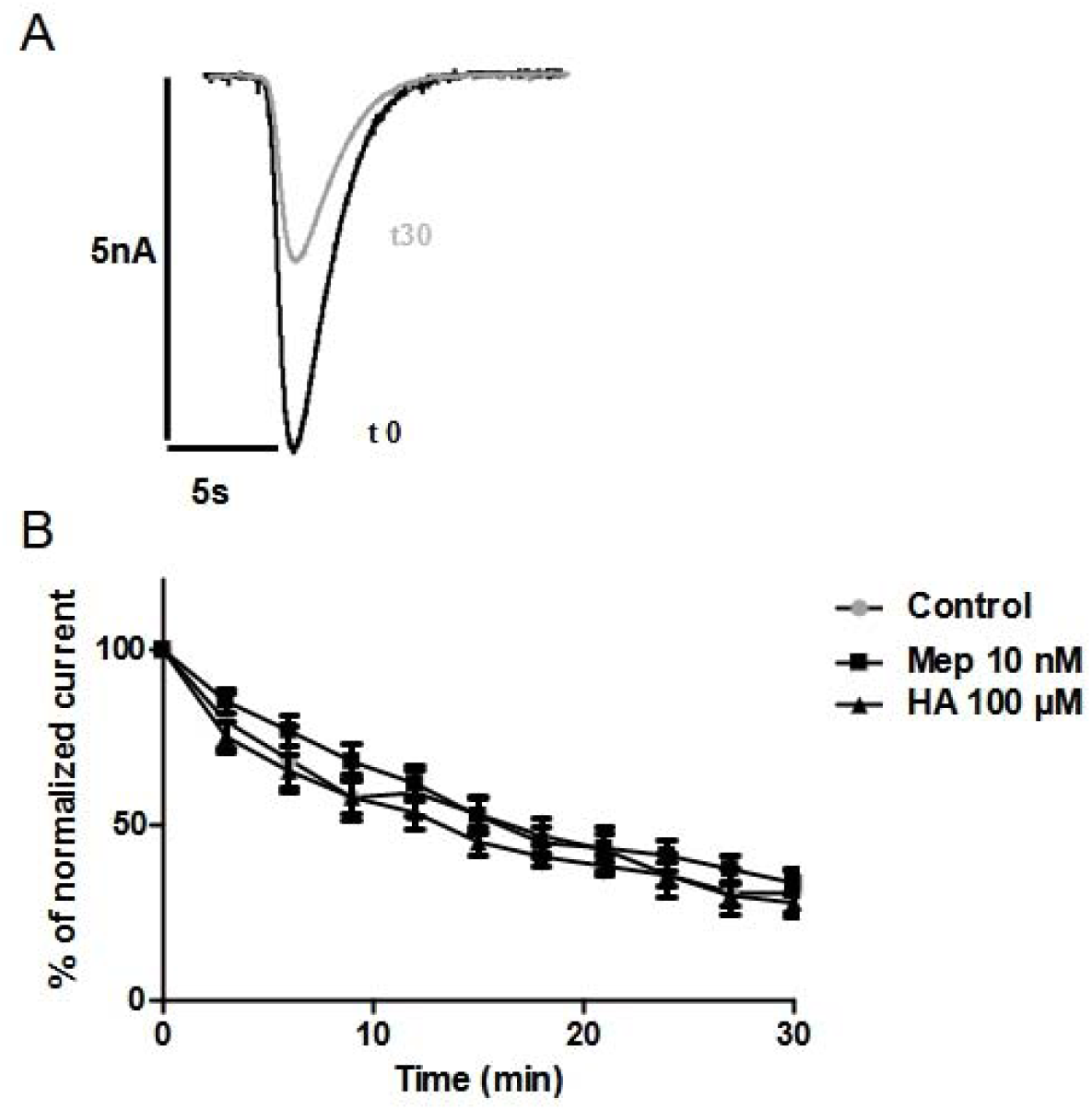
Effect of mepyramine (Mep) on the rundown of GABA currents. Rundown of GABA currents was observed by whole-cell recording. Inward currents were evoked by 1 s application of 100 µM GABA at 3 min intervals. **A** The rundown is shown by superposition of currents evoked after the first application (t0), and after 30 min (t30) of recording. **B** Currents, normalized to the first application (t0=100%), were evoked by application of GABA alone (n=12), with mepyramine 10 nM (n=10) or with histamine 100 µM (n=10).

### 3.6 Localisation of H_1_ receptors NMDA or GABA receptors

We first studied the localization of the H_1_ receptor immunoreactivity in the cultured hippocampal cells. The expression of the H_1_ receptor was observed within the vast majority (about 80 %) of the neurons (n=395 out of 500 HUC/ D positive cells) (Fig 6A).

**Figure 6.**
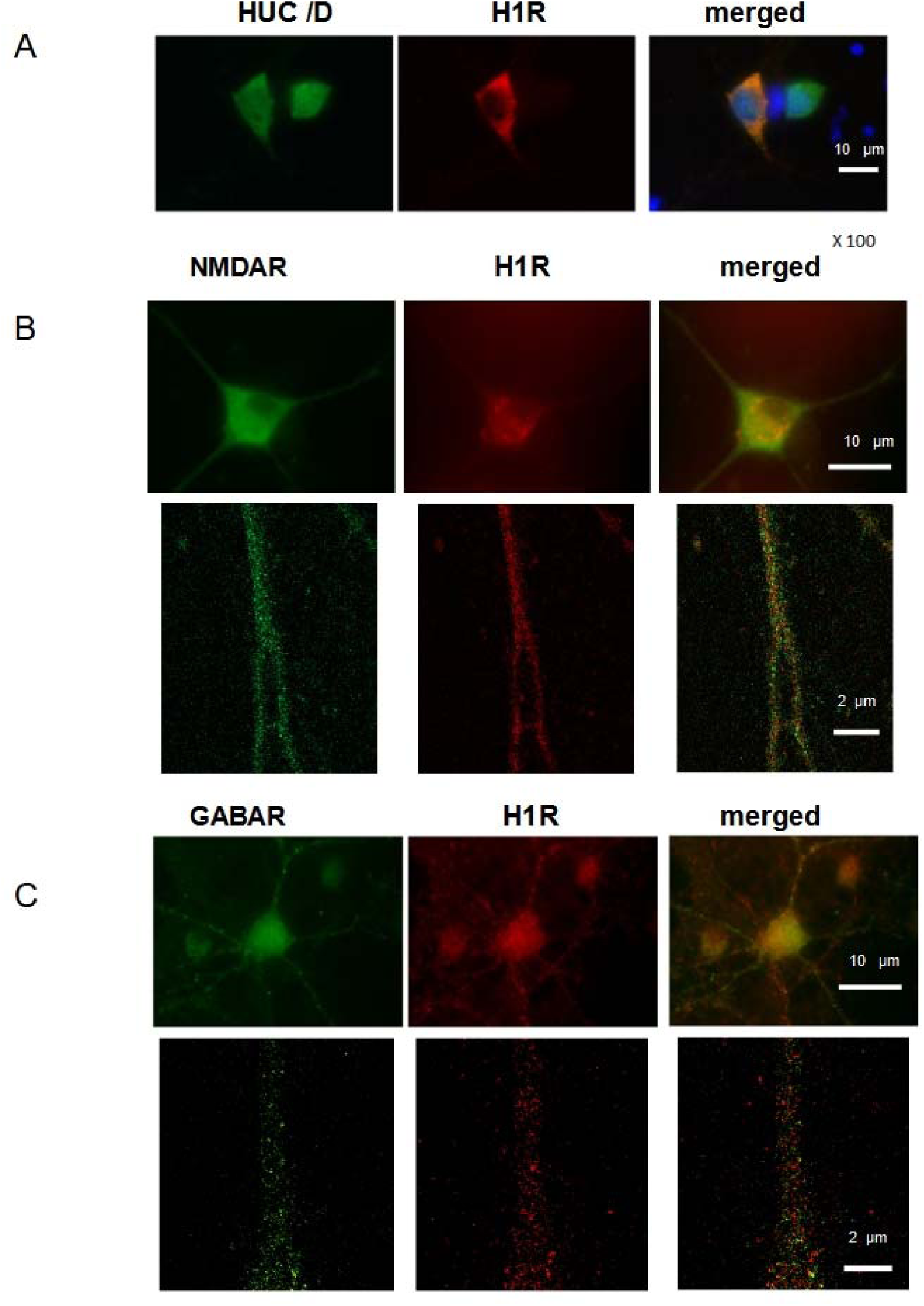
Immunostaining for the H_1_ receptor in neurons (HUC/D labeled cells) (A) in cultured hippocampal cells. Scale bar: 10 µm. Localisation of H_1_ receptors with NMDA receptors (B) and GABA_A_ receptors (C) in cultured hippocampal neurons, analysed at two different magnifications (scale bar: 10 µm or 2 µm, as indicated).

In a second step, the analysis at two different magnifications revealed that the H_1_ receptor immunoreactivity localised with the NMDA receptor (Fig 6B) as well as the GABA_A_ receptor (Fig 6C) in many neurons.

## 4 DISCUSSION

We used the rundown model to evaluate the effects of H1 receptor ligands, it is first important to note that in our experimental conditions we found the same results for our control curves, that Rosenmund (Rosenmund & Westbrook 1993), no difference in these curves with NMDA alone, so it seems that our experimental conditions do not modify the NMDA rundown. Similarly, by spacing the applications of NMDA to 3 minutes against 30 seconds during the recordings one avoids the appearance of the rundown NMDA and in these conditions we observed that the mepyramine did not affect the current NMDA, it is necessary to be in the condition of a fast repetitive application-inducing rundown to observe a modification of the NMDA answers. For the GABA recordings we have used exactly the same experimental conditions as Laschet (Laschet *et al* 2007) and no effect of mepyramine was observable.

This study reports for the first time that native histamine H_1_ receptors display constitutive activity. No endogenous histamine could be detected in the medium, evaluated with an enzyme immunoassay, indicating that mepyramine and triprolidine, two classical H_1_ antagonists, were acting here as inverse agonists, as previously observed with various antagonists at recombinant receptors (Bakker *et al* 2000; Fitzsimons *et al* 2004). Although very little data are available in rats, the potency of both inverse agonists in our study was in the same range as their potency previously reported at the guinea pig as well as human receptor (Bakkers et al 2001; Fitzsimons *et al* 2004). Our data are therefore consistent with a constitutive activity of the H_1_ receptor present in rat neuronal cultures, with mepyramine and triprolidine displaying the same nanomolar potency. Interestingly, triprolidine inhibited the constitutive activity of the guinea pig H_1_ receptor with a lower negative intrinsic efficacy than mepyramine, suggesting it might be classified as a partial inverse agonist, whereas mepyramine may be classified as a full inverse agonist (Fitzsimons *et al* 2004). This could explain the lower effect of triprolidine that we observe here compared to mepyramine used at the same concentration.

G protein-coupled receptors (GPCRs) are well known to be allosteric proteins that can adopt various conformations in equilibrium. A pharmacological entity such as protean agonism demonstrates the existence of constitutively active states of GPCRs different from and in competition with, ligand-directed active states (Gbahou *et al* 2003). This competition for G proteins leads to lower activation of the receptor by its agonists in the presence of a high degree of constitutive activity. In agreement with such a high constitutive activity of H_1_ receptors in our model, histamine itself and the more selective agonist 2-(3-bromophenyl)histamine (Leschke *et al* 1995) had no effect used alone. Consistent with our finding, no histamine-induced response could be detected when mutations of the H_1_ receptor resulted in a high constitutive activity (Bakker *et al* 2008). Moreover, the effect of both histamine and 2-(3-bromophenyl)histamine was restored when constitutive activity was decreased by mepyramine and triprolidine, i.e., when the equilibrium was shifted from constitutively active states towards ligand (agonist)-directed states. However, the observed effects of agonists against inverse agonists remained partial, which can be attributed to the fact that they promote different, i.e., active and inactive receptor conformations, respectively, leading their interaction to be more of an allosteric than a competitive and surmountable nature.

The second finding of our study shows that activation of H_1_ receptors does not modulate GABA receptors, but strongly modulates NMDA receptors. The increase of NMDA currents induced by H_1_ inverse agonists, and its reduction by H_1_ agonists, indicate that both constitutive activity and histamine decrease the activation of NMDA receptors. This effect, transient and short-lasting, clearly does not modulate the rundown of NMDA receptors itself, but requires the conditions of rundown to be observed, with none of the compounds displaying any effect after single administrations. This requirement strongly suggests that H_1_-receptor ligands do not interact directly with NMDA receptors. Histamine interacts with an allosteric site on NMDA receptors, but it induces a potentiation, and not a decrease, of NMDA currents. Moreover, H_1_-receptor ligands have no affinity for this site (Burban *et al* 2010).

Therefore, the delay induced in the settlement of the rundown is likely due to an interaction or crosstalk between the two receptors. The localisation of H_1_– and NMDA receptors that we observe in the majority of cultured hippocampal neurons show that this interaction may occur within the same neurons, although cell-cell interactions cannot be excluded. The presence of H_1_ receptors in cultures at embryonic day (E) 18 is consistent with their detection reported in the embryonic rat brain from the day (E) 14 (Kinnunen *et al* 1998). Further studies are required to identify which of the signaling pathways coupled to H_1_ receptors, are involved in the decrease of NMDA currents at initiation of rundown. H_1_-receptor activation has long been known to stimulate phospholipase C, even in the developing rat brain (Claro *et al* 1987). The resulting increase in intracellular (Ca^2+^) levels may delay the settlement of NMDA rundown (Rosenmund & Westbrook 1993). Interestingly, the reverse situation has been reported in the hippocampus, with NMDA inhibiting stimulation of phospholipase C by histamine (Baudry *et al* 1986). The effect, if any, of calcium on NMDA currents in our response is unclear but may be due to an increased density of NMDA receptors at the neuronal surface rather than an increase in the intrinsic activity of each entity. Although, not yet reported in neurons, H_1_-receptor activation increases intracellular (Ca2+) levels and reduces F-actin in cultured endothelial cells (Rotrosen & Gallin 1983). The calcium dependence of the NMDA rundown has been described in rat hippocampal neurons (Rosenmund & Westbrook 1993) and actin depolymerization induced by calcium reduces NMDA channel activity by altering the interaction between NMDA receptors and actin filaments (Rosenmund & Westbrook 1993’). Such crosstalk between NMDA and H_1_ receptors via intracellular (Ca2+) and/or actin is consistent with a modulation of membrane targeting of NMDA receptors by H_1_-receptor activation. Interestingly, a rapid surface accumulation of NMDA receptors increases glutamatergic excitation during status epilepticus (Naylor *et al* 2013), and an increase of H_1_-receptor densities is observed in the brain of rats with generalized epilepsies (Midzyanovskaya *et al* 2016).

An alternative mechanism is based on the links between metabolic signaling and neuronal excitability. In early epileptogenesis, metabolic reprogramming triggers inhibition of AMPK (AMP-Activated Protein Kinase) and subsequent activation of mTOR, which is a signature event of epileptogenesis (Alqurashi *et al* 2021). Neuronal plasticity, including LTP, is thought to play an important role in the progression of seizures to intractable epilepsy and AMPK plays an important role as an energy sensor in the metabolic regulation of neuronal plasticity and its aberrant forms such as epilepsy (Potter *et al* 2010). Interestingly, weight gain induced by some antipsychotic drugs is mediated by histamine H_1_ receptor-linked activation of hypothalamic AMP-kinase (Kim *et al* 2007). In addition, although they were used at a very high concentration, various H_1_-receptor inverse agonists, such as triprolidine, potentiate responses, oxygen radical formation, and excitotoxicity induced by NMDA in cerebellar neurons, effects that were prevented by the addition of histamine (Díaz-Trelles *et al* 2000).

Whatever the mechanism involved, our data may underlie, at least partially, the well-established suppressive role of histamine in seizure development through H_1_-receptors, as well as its counterpart, i.e.,the epileptogenic activity of (brain penetrating) first-generation H_1_ antagonists/inverse agonists. Activation of H_1_ receptors by constitutive activity or histamine itself would tend to counteract the excessive NMDA receptor activity, and therefore an enhanced susceptibility to status epilepticus.

The modulation of NMDA receptors by H_1_ receptors may also explain, at least partially, the well-known higher susceptibility of children to status epilepticus, compared to adults. The first generation of H_1_-receptor inverse agonists, used as classical anti-allergy drugs, was reported to induce occasional convulsions in healthy children as well as patients with epilepsy (Churchill 1949; Yasuhara *et al* 1998; Yokoyama & Iinuma 1996). In childhood, this higher susceptibility remains poorly understood, and has been globally related to the immaturity of the developing brain (Yokoyama 2001). In electrically-induced seizures, various brain-penetrating H_1_-receptor antagonists/inverse agonists, including mepyramine, increase the durations of all the convulsive phases in young mice but not in adult mice (Yokoyama *et al* 1993). Interestingly, H_1_-receptor knock-out mice also show age-dependent susceptibility to status epilepticus. More severe seizures, compared to those of wild type mice, were reported in immature mice (Kukko-Lukjanov *et al* 2010), but not in adult mice (Kukko-Lukjanov *et al* 2012). Moreover, it is worth noting that studies based on neuronal cultures inherently deal with the developing brain. Consistent with our findings, histaminergic neurons protect the developing hippocampus from kainic acid-induced neuronal damage in an organotypic coculture system (Kukko-Lukjanov *et al* 2006). Altogether, these observations would suggest that the modulation that we report here is less operatin in normal adults. However, further studies are needed to know if the high degree of constitutive activity is still present in the normal adult brain, or if the activation of H_1_-receptor by histamine becomes progressively predominant during aging. Consistent with such an evolution, the administration of a selective H_1_ agonist decreases pentylenetetrazole-induced convulsions in adult mice, an effect antagonized by centrally acting H_1_ antagonists such as mepyramine (Yokoyama *et al* 1994). At last, blockade of histamine H_3_ autoreceptors, known to increase the release of endogenous histamine in the brain, reduced photosensitivity in adult epileptic patients (Kasteleijn-Nolst *et al* 2013) and inhibited pentylenetetrazole-kindled seizures in adult rats, an effect reversed by administration of mepyramine (Zhang *et al* 2003).

## Conclusion/Summary

◦ Histamine has a well-established suppressive role in seizure development via activation of H_1_-receptors.
◦ Brain penetrating H_1_ inverse agonists display an epileptogenic activity.
◦ Two H_1_ inverse agonists increase NMDA currents and delay the initiation and settlement of NMDA rundown in neuronal cultures, an effect reversed by histamine and a selective H_1_ agonist.
◦ During the initiation of rundown, the increase induced by inverse agonists reveals a strong modulation of NMDA receptors by the constitutive activity of H_1_ receptors and histamine, which may underlie experimental and clinical observations.

## Abbreviations

AUC: area under the curve
GPCRs: G protein-coupled receptors
Mep: mepyramine
Tri: triprolidine
HA: histamine
Bro: 2,3 bromophenylhistamine

## Acknowledgements

This study was supported by INSERM, the French Ministère de la Recherche.

## Author contributions

**E**xperiments were designed and planned by VA and JMA. Electrophysiology, immunohistochemistry experiments and analysis were carried VA, VA and JMA wrote the manuscript. All authors have approved the final version of the manuscript and agree to be accountable for all aspects of the work. All persons designated as authors qualify for authorship, and all those who qualify for authorship are listed.

